# Generation of a global freshwater algal taxonomic database by application of PCR-free *rbcL* gene detection and machine-learning-based taxonomic classification to public metagenome datasets

**DOI:** 10.1101/2025.08.28.672922

**Authors:** Robert W. Murdoch

## Abstract

While it has become increasingly evident that microalgae are critical components of aquatic ecosystems, comprehensive taxonomic analysis of the microalgal component of microbial communities has lagged behind recent innovations in prokaryotic sequencing and bioinformatics. Microalgae are sequenced alongside prokaryotes in metagenomic whole genome shotgun (WGS) sequencing efforts but remain overlooked. In this study, an analytic pipeline for detecting a key algal taxonomic barcode, *rbcL*, from WGS datasets. This approach allowed for near-perfect detection of *rbcL* sequences from eight major algal phyla, whereas *in silico* PCR utilizing popular phyla-specific *rbcL* primer sets for four algal phyla suggests that such approaches may miss much algal diversity. Further, a machine learning classification model was built and validated to allow for taxonomic classification of *rbcL* sequences of eight algal phyla to genus or species level with high accuracy and recall values (>90%). Benchmarking of this approach against microscopic algal identifications showed accurate identifications for algae with higher relative abundance values and may offer additional detections of algae not readily identified by microscopy. This pipeline was applied against over 4,000 freshwater WGS metagenomes to generate a global taxonomic occurrence and relative abundance database of freshwater algae composed of over 14,000 accessions. WGS-based algal detection offers broad, unbiased taxon-agnostic algal detection in a single sample analysis. Furthermore, it allows for simultaneous identification of the prokaryotic community, offering new opportunities for studying inter-domain relationships as evidenced by a case-study presented herein. The *rbcL* taxonomic classifier and metagenome-derived algal detection database are available for download and use at https://doi.org/10.6084/m9.figshare.29996962.

## Introduction

Microalgae are important members of the global ecosystem, responsible for roughly half of global carbon fixation ^1^ and serving as the foundation of aquatic food webs around the world. The polyphyletic nature of algae ^2^, presence of diverse taxonomic barcode genes, and paucity of available genomic resources have limited holistic, pan-phyletic identification and enumeration of algal-prokaryotic communities via high-throughput sequencing methods ^3^. Given the absence of agnostic tools, microalgal researchers continue to refine and apply group-specific primer sets for PCR amplicon library sequencing. While this has opened the ability to identify multiple algal taxa within a selected phylum simultaneously, the reliance on PCR amplification introduces considerable restraints and pitfalls; because the most taxonomically informative barcodes amongst many algal taxa are protein-coding genes (ex. *rbcL*) rather than rRNA genes (reviewed in ^4, 5^), nucleotide sequences among taxa vary to such a degree that universal primer design has proven infeasible. While there are broad-specificity degenerate primer sets available for certain algal taxa (ex. brown algae ^6^, green algae ^7^, and diatoms ^8^), many taxa remain recalcitrant to such study. Additionally, PCR amplification using such degenerate primer sequences is known to be prone to potentially drastic inter-taxa biases, distorting the true state of the underlying community. Furthermore, focus on PCR of algal barcodes excludes prokaryotes entirely. Research to date indicates that microalgae extensively interact with prokaryotes via resource competition, chemical antagonism, and nutrient exchanges ^9^, interactions with emergent consequences such as harmful algal blooms and methane production ^10^.

Whole-genome shotgun (WGS) metagenome sequencing fundamentally allows for simultaneous insights into all genetic material present in a community DNA sample. While still prone to biases in extraction and sequencing efficiency among taxa, WGS avoids community pattern distortions that arise due to taxonomy-driven PCR amplification blind-spots and biases ^11^. PCR bias arises due to differential binding affinity inherent in the reliance on degenerate primer sequences and imperfect template matching that can distort true template relative abundances dramatically (reviewed in ^12^). Additionally, primer design is by necessity limited by availability of reference sequences and thus may perform poorly or not at all against microalgal clades with sparse reference sequences to design against. Environmental microbiology continues to discover novel clades of unculturable prokaryotes via WGS metagenomics, clades that had previously been undiscovered because “universal” 16S rRNA primers did not account for their unique genetic characteristics ^13^. The rapidly decreasing costs of WGS sequencing vs. amplicon libraries have driven widespread adoption of WGS for microbial community analysis.

While over 100,000 WGS metagenomes have been assembled and released into public databases ^14^, the potential microalgal components of these systems have been overlooked due to lack of accessible, validated analytic pipelines. While metagenomes are typically generated for study of prokaryotic components, microalgae may be sequenced alongside them yet simply ignored. To our knowledge, few current metagenome analytic pipelines consider microalgae - whole-genome-similarity based pipelines suffer from the general paucity of available algal genomes ^15^, while pipelines that recruit and classify 18S rRNA reads ^16^ will produce lower accuracy taxonomic assignments. Microalgal taxonomic barcodes such as *rbcL* are likely sequenced alongside prokaryotic components and if so, can be detected and classified without the need for PCR amplification.

In this study, an approach for harvesting *rbcL* sequences from metagenomes was developed and applied to all 4,206 freshwater metagenomes available in the JGI Integrated Microbial Genomes (IMG) database (available as of February 2023) to 1) leverage large quantities of publicly available sequencing resources and 2) lay the groundwork for a novel approach for characterizing microalgae in complex communities. Additionally, due to lack of established state-of-the-art tools for sequence classification of *rbcL* sequences, an AI/ML taxonomic classifier was trained, validated, and applied to harvested *rbcL* genes. AI/ML classification of eukaryotic barcodes have shown promise for plants ^17^ but have yet to be applied to algae. Performance of the tool against microscopy-driven taxonomic assessment is assessed, and a case study is presented to demonstrate holistic study of the prokaryotic and algal communities.

## Methods

### rbcL taxonomic classifier training and validation

All *rbcL* accessions were collected from the Barcode of Life Database (BOLD) Feb 4, 2023 public release ^18^. Bacterial sequences and accessions without species assignments were removed, leaving 91,997 sequences. High-confidence and informative bacterial *rbcL* sequences were amended to the database. These included all *rbcL* collected from isolate genomes in the JGI IMG database ^19^ corresponding to phylum Cyanobacteriota and all American Type Culture Collection (ATCC) strains, 565 in total.

The single longest sequence for each species was utilized for classifier training (43,315 sequences) while the remaining 49,247 sequences were utilized for model validation. Training set sequences and their corresponding seven-level taxonomy were used to train a naïve-Bayes classifier operated in a qiime2-amplicon-2023.9 conda environment ^20^. The classifier was operated under default settings, which employs a lower confidence threshold of 0.7. The validation sequences were then ingested, classified, and resulting classifications compared to the “known” classifications provided by the BOLD database. Coverage and accuracy rates for each phylum at each taxonomic sub-level and across confidence cut-off values of 0.7-0.99 were independently calculated by equations A and B ^21^.

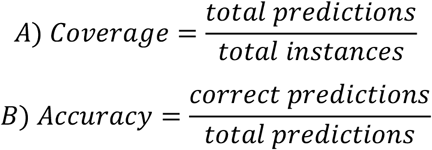

### Detection and taxonomic classification of rbcL genes in metagenomes

The efficacy of the pfam00016 ribulose bisphosphate carboxylase large chain, catalytic domain (*i*.*e. rbcL*) annotation for detecting *rbcL* genes was assessed by searching BOLD *rbcL* accessions for its presence ^22^. To restrict pfam00016 efficacy analysis only to nearly-complete sequences and reduce taxonomic sampling bias, the database was curated to retain only sequences above 1,200bp and only one per species. The resulting 28,067 *rbcL* sequences were translated into most likely protein sequences by FragGeneScan v1.2 ^23^. FragGeneScan was operated using the “sanger_5” training model, which assumes up to 0.5% sequencing error. The resulting protein sequences were scanned against the pf00016 hmm model using the hmmscan command in HMMER 3.1b2 under default settings ^24^. Domain gathering bit-score threshold was established as 30.1, reflecting the model-specific cut-off recommendation that is also applied by the IMG annotation pipeline.

All 4,206 assembled freshwater metagenomes in the IMG database, defined as those with the metadata tag “Ecosystem Type = Freshwater”, were delineated on February 23, 2023 and scanned for genes that had been assigned domain annotation pfam00016 by the IMG metagenome annotation pipeline. Gene nucleotide sequences were downloaded along with source genome metadata and estimated scaffold read depth metrics. Nucleotide sequences and metagenome details for the resulting 150,811 sequences are provided in Supplementary Table S1. Genes were classified using the constructed *rbcL* taxonomic classifier following import into qiime2-amplicon-2023.9 operated with a minimum confidence threshold of 0.7.

### Simulated PCR to assess primer efficacy against metagenome detected rbcL sequences

The efficacy of several current algal primer sets against metagenome-derived microalgal *rbcL sequences* was estimated by *in silico* PCR. The primer sets applied are detailed in Table 1. PCR simulation was conducted using thermonucleotideBLAST v2.66 ^25^ operated using default parameters and a permissive primer annealing temperature of 40C. To account for the potential lack of primer annealing sites presented by fragmentary *rbcL* sequences, only genes 1,200 bp and longer were included in analyses.

**Table 1.**
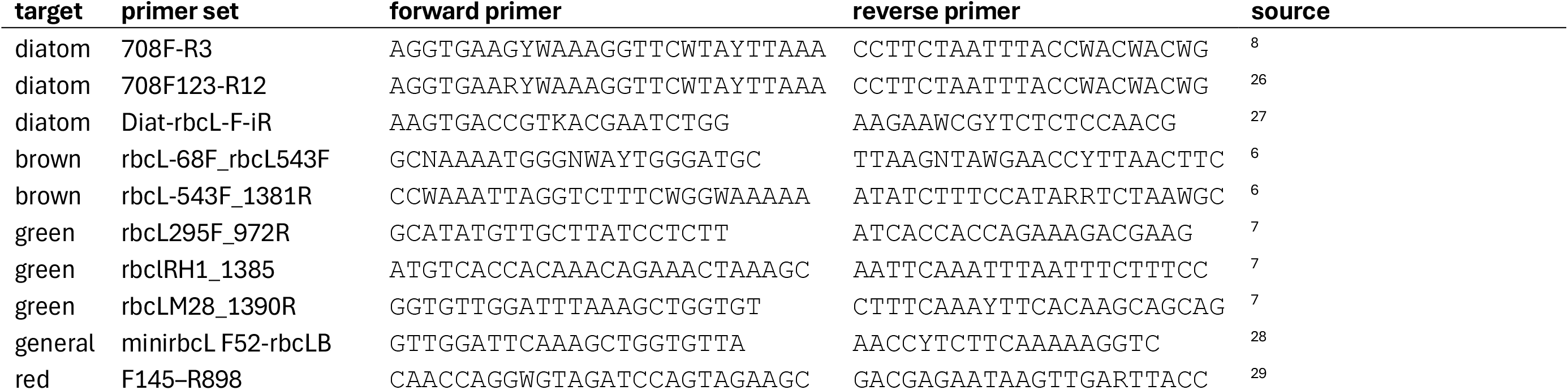
Primer sets tested by *in silico* PCR against high-quality BOLD *rbcL* gene sequences.

### Benchmarking against microscopically classified algal communities

Microscopically classified diatom and soft algae sample data was obtained from the National Science Foundation National Ecological Observatory (NSF NEON) data repository ^30^ and compared to algal identifications derived from WGS metagenomic *rbcL* taxonomic classification. All six 2018 “periphyton, seston, and phytoplankton collection” algal identification data for West St. Louis Creek, Colorado (WLOU) NEON was downloaded and binned. The samples were comprised of a mixture of seston, epilithic, and epipsammic samples. Non-algal classifications (bacterial and charophyte) were removed and relative abundances calculated as mean per bottle cell counts averaged across samples. Corresponding 2018 benthic WGS data from the WLOU site, NEON data product “Benthic microbe community composition”, were downloaded from the NEON data portal. Because data at that year were only provided as unassembled raw sequencing reads, data was subjected to local quality control (QC), assembly, gene calling, and *rbcL* domain detection. QC was conducted using fastp v0.23.1, operated so as to preserve paired reads or merge read pairs in the case of overlap as appropriate ^31^. Metagenome assemblies were generated using MEGAHIT v1.2.9 ^32^ utilizing all read pairs and corresponding merged sequences representing each collection date. MEGAHIT was operated using the “meta-large” preset. Genes and their protein products were predicted resulting six assemblies were using Prodigal v2.6.3 operated with the “-meta” flag ^33^. *rbcL* domains in the resulting protein sequences were detected using HMMER 3.1b2 as described above. In addition, *rbcL* genes in a set of eight 2023 WGS metagenomes hosted at IMG representing the same WLOU NEON site (IMG genome IDs 3300080923, 3300080924, 3300081714, 3300081902, 3300081903, and 3300081911 – 3300081913) were harvested and subjected to taxonomic classification as described above.

### Calculating total microbial and algal relative abundance in metagenomes

Total microbial gene copy number per metagenome was estimated by summing the read depth of the single copy conserved protein (SCCP) gene encoding ribosomal protein L2 (COG0090). GTDB-derived taxonomy ^34^ of the scaffold harboring the L2 gene, provided by the IMG database, was used to estimate prokaryotic taxonomy.

## Results and Discussion

### Taxonomic Classifier Model Validation and Performance

Training and validation sequences were abundant (25 or more training sequences) for four major algal phyla, ten plant phyla, and Cyanobacteriota (Table 2). It should be noted that the taxonomy employed in this study by necessity reflects that used by BOLD, wherein the diatoms are placed at phylum level (Bacilliarophyta). Examination of model performance vs. confidence indicated at increasing the confidence threshold led to significant drops in coverage with only minor increases in accuracy (Figure 1). While different application spaces might warrant different coverage vs. accuracy trade-offs, the minimum confidence score of 0.7 was chosen for further examination of model performance, benchmarking, case-study, and database generation.

**Table 2.**
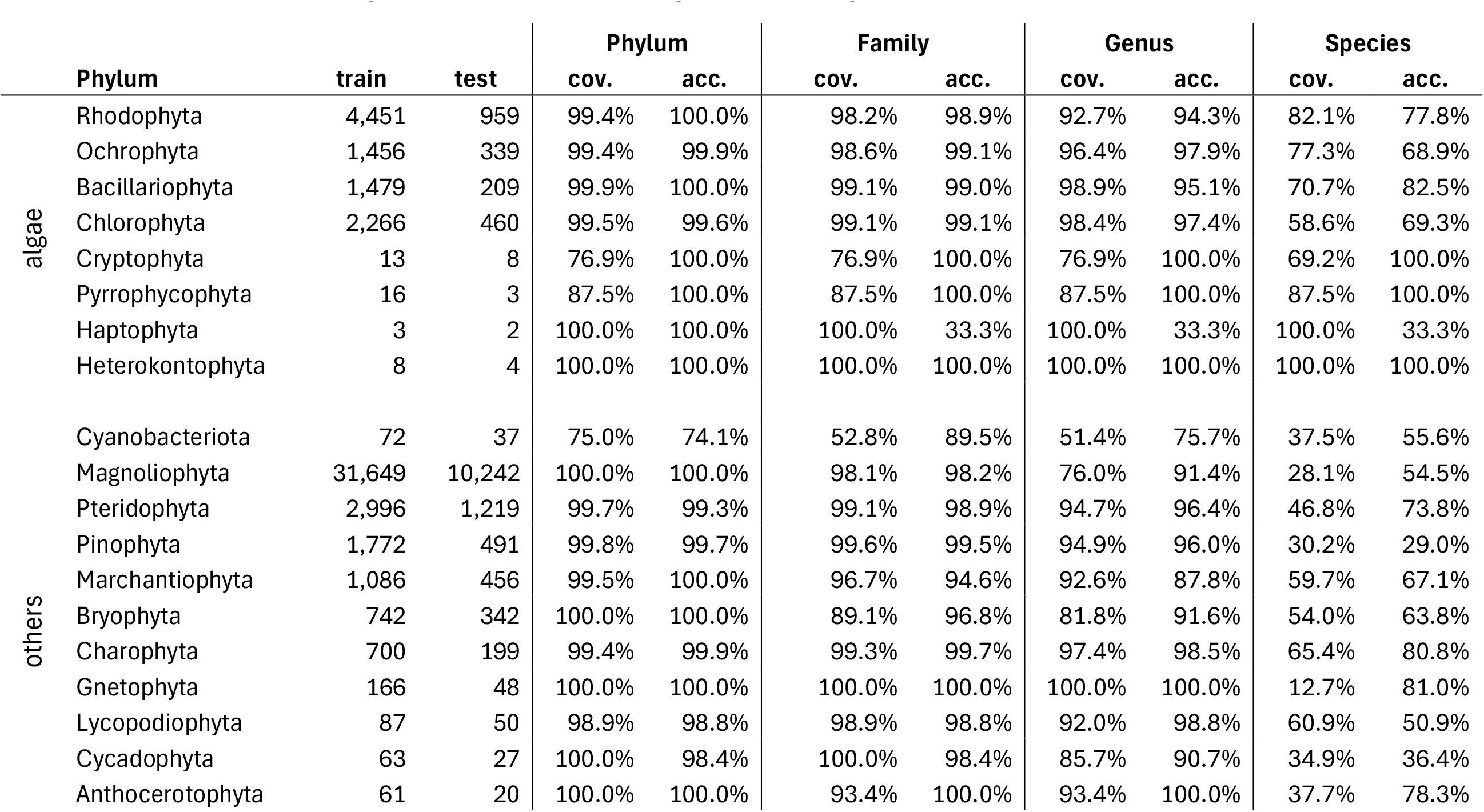
Taxonomic classifier model performance across phyla. “train” indicates the number of reference sequences from the indicated phylum used to train the classifier, while “test” indicates the number of sequences on which model performance was validated. Model performance validation at the indicated taxonomic level is quantified as the percentage of test sequences which were assigned a taxonomic classification (coverage, “cov.”) and the percentage of those assignments which were correct (accuracy, “acc.”).

|  | Phylum | train | test | Phylum |  | Family |  | Genus |  | Species |  |
| --- | --- | --- | --- | --- | --- | --- | --- | --- | --- | --- | --- |
|  |  |  |  | cov. | acc. | cov. | acc. | cov. | acc. | cov. | acc. |
| algae | Rhodophyta | 4,451 | 959 | 99.4% | 100.0% | 98.2% | 98.9% | 92.7% | 94.3% | 82.1% | 77.8% |
|  | Ochrophyta | 1,456 | 339 | 99.4% | 99.9% | 98.6% | 99.1% | 96.4% | 97.9% | 77.3% | 68.9% |
|  | Bacillariophyta | 1,479 | 209 | 99.9% | 100.0% | 99.1% | 99.0% | 98.9% | 95.1% | 70.7% | 82.5% |
|  | Chlorophyta | 2,266 | 460 | 99.5% | 99.6% | 99.1% | 99.1% | 98.4% | 97.4% | 58.6% | 69.3% |
|  | Cryptophyta | 13 | 8 | 76.9% | 100.0% | 76.9% | 100.0% | 76.9% | 100.0% | 69.2% | 100.0% |
|  | Pyrrophytophyta | 16 | 3 | 87.5% | 100.0% | 87.5% | 100.0% | 87.5% | 100.0% | 87.5% | 100.0% |
|  | Haptophyta | 3 | 2 | 100.0% | 100.0% | 100.0% | 33.3% | 100.0% | 33.3% | 100.0% | 33.3% |
|  | Heterokontophyta | 8 | 4 | 100.0% | 100.0% | 100.0% | 100.0% | 100.0% | 100.0% | 100.0% | 100.0% |
| others | Cyanobacteriota | 72 | 37 | 75.0% | 74.1% | 52.8% | 89.5% | 51.4% | 75.7% | 37.5% | 55.6% |
|  | Magnoliophyta | 31,649 | 10,242 | 100.0% | 100.0% | 98.1% | 98.2% | 76.0% | 91.4% | 28.1% | 54.5% |
|  | Pteridophyta | 2,996 | 1,219 | 99.7% | 99.3% | 99.1% | 98.9% | 94.7% | 96.4% | 46.8% | 73.8% |
|  | Pinophyta | 1,772 | 491 | 99.8% | 99.7% | 99.6% | 99.5% | 94.9% | 96.0% | 30.2% | 29.0% |
|  | Marchantiophyta | 1,086 | 456 | 99.5% | 100.0% | 96.7% | 94.6% | 92.6% | 87.8% | 59.7% | 67.1% |
|  | Bryophyta | 742 | 342 | 100.0% | 100.0% | 89.1% | 96.8% | 81.8% | 91.6% | 54.0% | 63.8% |
|  | Charophyta | 700 | 199 | 99.4% | 99.9% | 99.3% | 99.7% | 97.4% | 98.5% | 65.4% | 80.8% |
|  | Gnetophyta | 166 | 48 | 100.0% | 100.0% | 100.0% | 100.0% | 100.0% | 100.0% | 12.7% | 81.0% |
|  | Lycopodiophyta | 87 | 50 | 98.9% | 98.8% | 98.9% | 98.8% | 92.0% | 98.8% | 60.9% | 50.9% |
|  | Cycadophyta | 63 | 27 | 100.0% | 98.4% | 100.0% | 98.4% | 85.7% | 90.7% | 34.9% | 36.4% |
|  | Anthocerotophyta | 61 | 20 | 100.0% | 100.0% | 93.4% | 100.0% | 93.4% | 100.0% | 37.7% | 78.3% |

**Figure 1.**
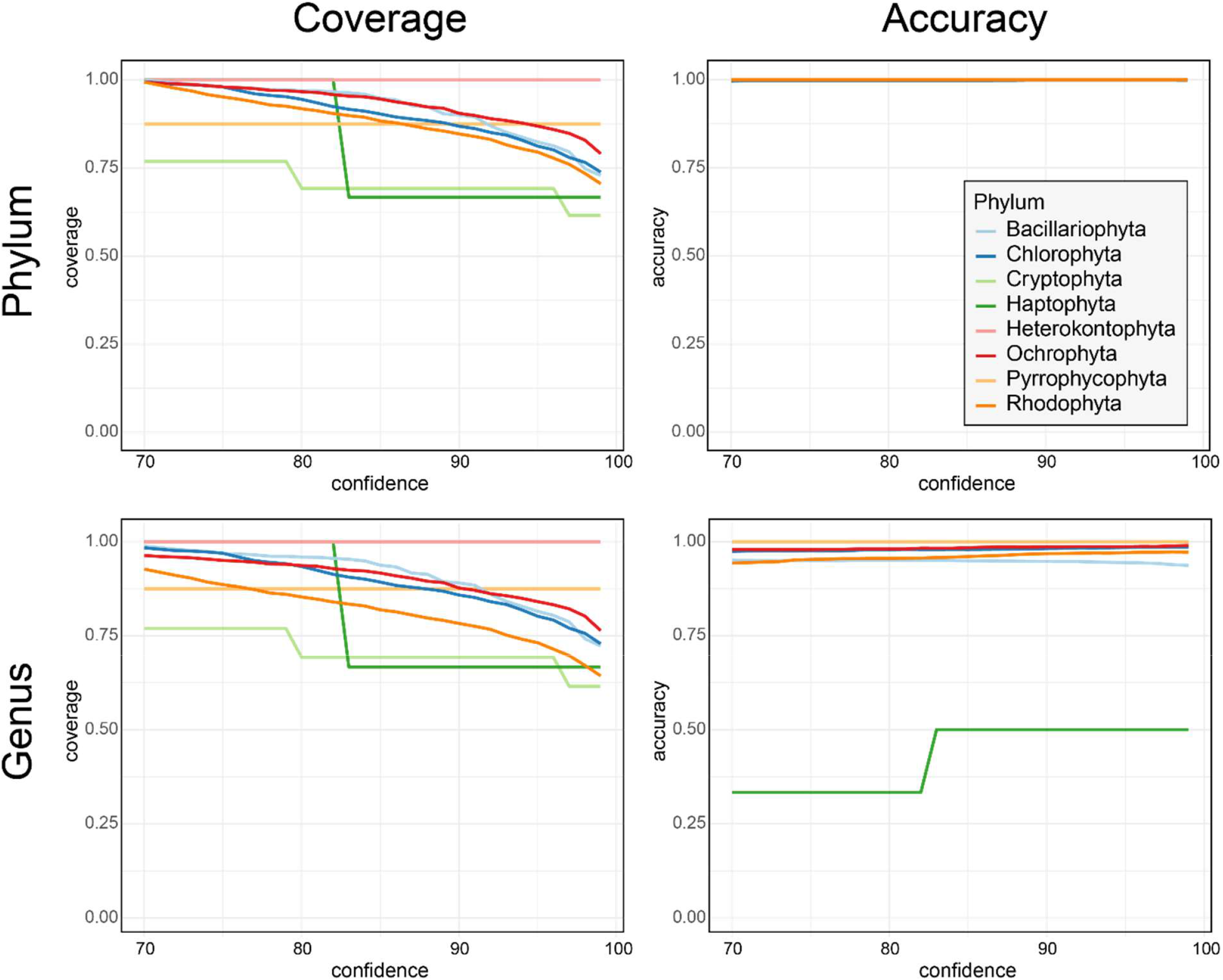
Model performance vs. confidence threshold. Coverage and accuracy percentages (y-axes) of classifier phylum and genus assignments across confidence threshold cut-offs ranging from 70% to 100% (x-axis).

Model validation indicated near-perfect (>98% coverage and accuracy) family-level resolution of the four most informed algal phyla with the most *rbcL* reference sequences – Chlorophyta, Bacillariophyta, Ochrophyta, and Rhodophyta. Genus-level classification coverage and accuracy remained above 92% at genus level but dropped to 69%-89% accuracy for species-level classifications (Table 2). This is consistent with current practices whereby the *rbcL* is generally regarded as an informative barcode for these taxa ^5^. The other four algal phyla, Haptophyta, Heterokontophyta, Cryptophyta, and Pyrrophycophyta, had low available sequences for training and validation of the taxonomic classifier. However, the validation results indicated successful classification (>75% coverage and 100% accuracy at phylum level for all four and same metrics down to family level excluding Haptophyta). The paucity of *rbcL* sequences in the BOLD database identified down to species level for these clades suggests caution in interpretation of sample analytic results indicating these taxa.

The non-Chlorophyta plant classifications were near-perfect down to family level, although performance began to drop at genus and species level (Table 1), consistent with many reports that *rbcL* is informative but not comprehensive for land plant taxonomy ^35^. The model generally performed poorly for cyanobacteria, showing only ∼75% coverage and accuracy at Phylum level and 51% / 76% coverage/accuracy at genus level – this lack of performance was consistent with known limited taxonomic utility of *rbcL* among Cyanobacteria due to lateral gene transfer ^36^. Cyanobacteria and non-chlorophyte plant delineation by *rbcL* is not considered further in this report.

The trained qiime2-amplicon-2023.9 *rbcL* taxonomic classifier is available in the supplementary materials.

### Algal rbcL classification allows for robust identification of algae in freshwater metagenomes

The pfam00016 rbcL domain was detected in 99.6% of the 28,067 representative BOLD *rbcL* sequences, confirming the utility of this pfam domain for *rbcL* delineation. 150,811 *rbcL* genes and gene-fragments were detected in 2,611 of the 4,206 freshwater metagenomes analyzed. The metagenomes in which *rbcL* sequences were detected originated from around the world, although with higher density in North America and Europe (Figure 2).

**Figure 2.**
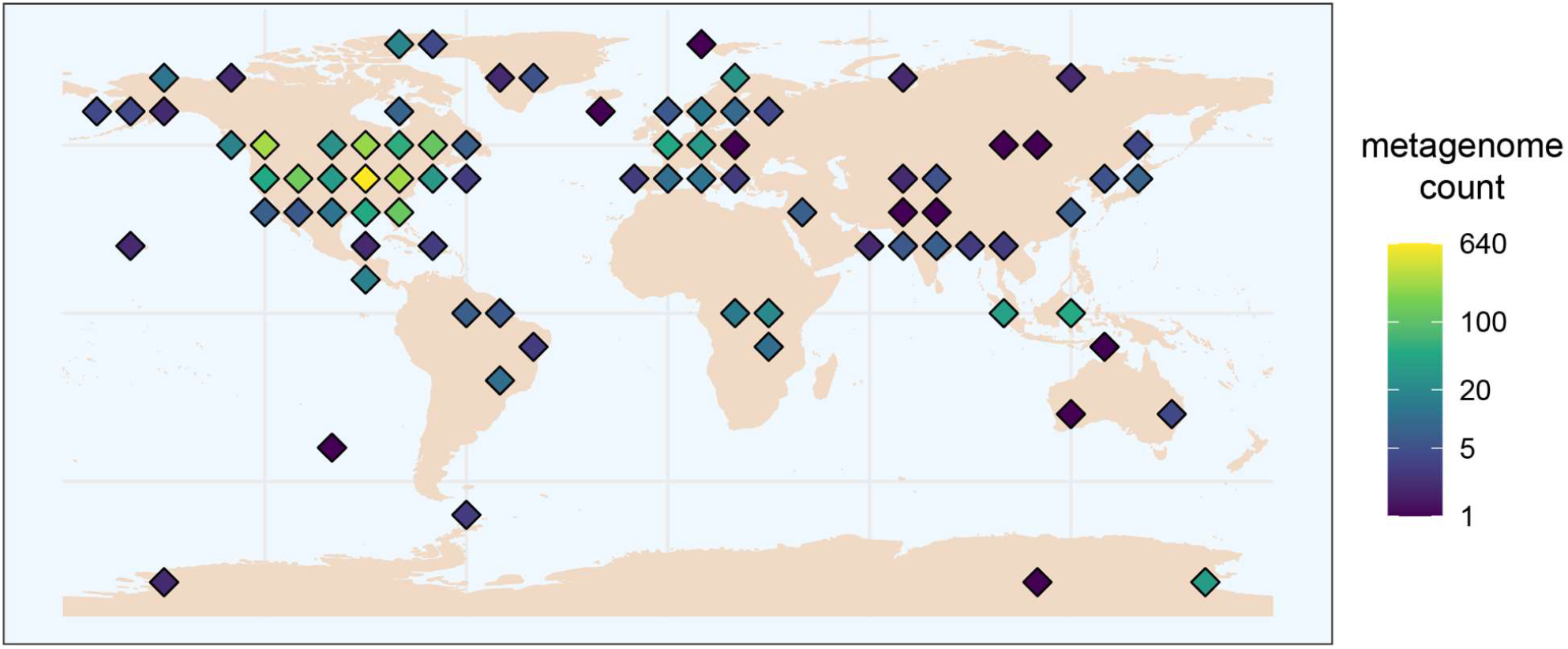
Heatmap where color corresponds to number of freshwater metagenomes with detected *rbcL* sequences. Sample geocoordinates are rounded to the nearest 10 degrees latitude and longitude.

All *rbcL* gene sequences detected in IMG freshwater metagenomes were downloaded and taxonomically classified. The sequences were generally fragmentary, with an average length of 625 bp, although 19,056 were complete or nearly complete (>1,200bp). A total of 14,931 sequences were classified as one of the eight trained algal taxa or as unassigned members of domain Protista, which may represent other *rbcL-*harboring protists not represented in the database (Table 3). The most commonly detected phyla were Bacillariophyta, Ochrophyta, and Chlorophyta, consistent with common conception of freshwater algal communities. The full database of metagenomic *rbcL* sequences with corresponding IMG gene and genome IDs, sequence taxonomic classification and confidence scores, and metagenome metadata are available in Supplementary Table S1 (all *rbcL* sequences) and Supplementary Table S2 (only algal *rbcL* sequences).

**Table 3.** Total numbers of freshwater metagenome-derived *rbcL* taxonomically classified as the indicated phylum.

| kingdom | phylum | n |
| --- | --- | --- |
| Plantae | Chlorophyta | 2,934 |
| Protista | Bacillariophyta | 4,143 |
| Protista | Cryptophyta | 1,837 |
| Protista | Haptophyta | 451 |
| Protista | Heterokontophyta | 13 |
| Protista | Ochrophyta | 3,350 |
| Protista | Pyrrophytophyta | 682 |
| Protista | Rhodophyta | 51 |
| Protista | *unassigned* | 930 |

Copy-number weighted phyla comparisons, reflecting relative abundance of each sequence among all algae detected per sample, largely reflected unique sequence counts (Table 4) – Bacillariophyta comprised 40.1% of all copy-number-weighted sequences, followed by Ochrophyta and Chlorophyta, with 20.5% and 14.6% respectively. These three phyla collectively dominated most freshwater ecosystems. The Cryptophyta, Pyrrophycophyta (dinoflagellates), and Haptophyta also comprised significant proportions of copy-number-weighted sequences (14.2%, 6.9%, and 3.4% respectively), making up notable proportions of lake and ice communities. Heterokontophyta and Rhodophyta were only rarely detected, primarily occurring in pond and creek ecosystems, respectively.

**Table 4.**
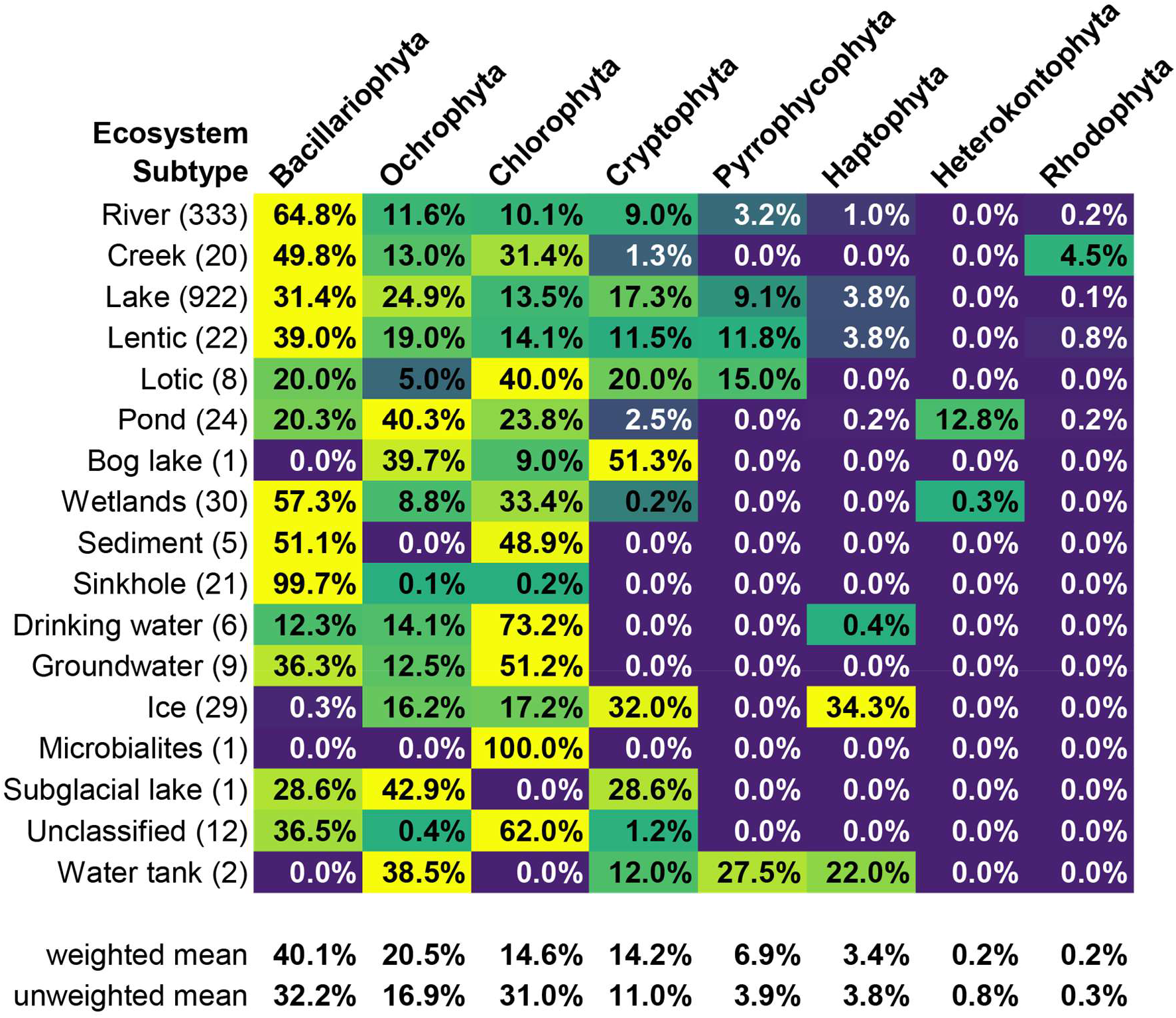
Read depth weighted relative abundances of algal phyla by freshwater ecosystem type.

### PCR-based approaches may miss much true algal diversity

Compared to PCR amplicon sequencing, WGS metagenome analysis offers the distinct advantage of comprehensive algal detection in a single sample analysis. Established *rbcL* primers are not even available for four major algal clades (Cryptophyta, Haptophyta, Heterokontophyta, and Pyrrophycophyta), although primers for alternative barcodes have been developed ^37^. An *in silico* assessment of primer sets that target *rbcL* of four major algal clades (Chlorophyta, Bacilliarophyta, Ochrophyta, and Rhodophyta) suggested variable efficacy against high-quality (1200+ bp) *rbcL* sequences that were detected using the conserved-domain-based (pfam00016) metagenome strategy (Table 5). Bacilliarophyta primers performed well, amplifying up to 933 of 942 available Bacilliarophyta sequences (99.0%), consistent with the widespread consensus that *rbcL* is an excellent and accessible barcode for the diatoms ^8^. Primers targeting the other three clades performed less well, amplifying at best 54.5% of Chlorophyta, 79.7% of Ochrophyta, and 1 of 5 Rhodophyta sequences (Table 5).

**Table 5.**
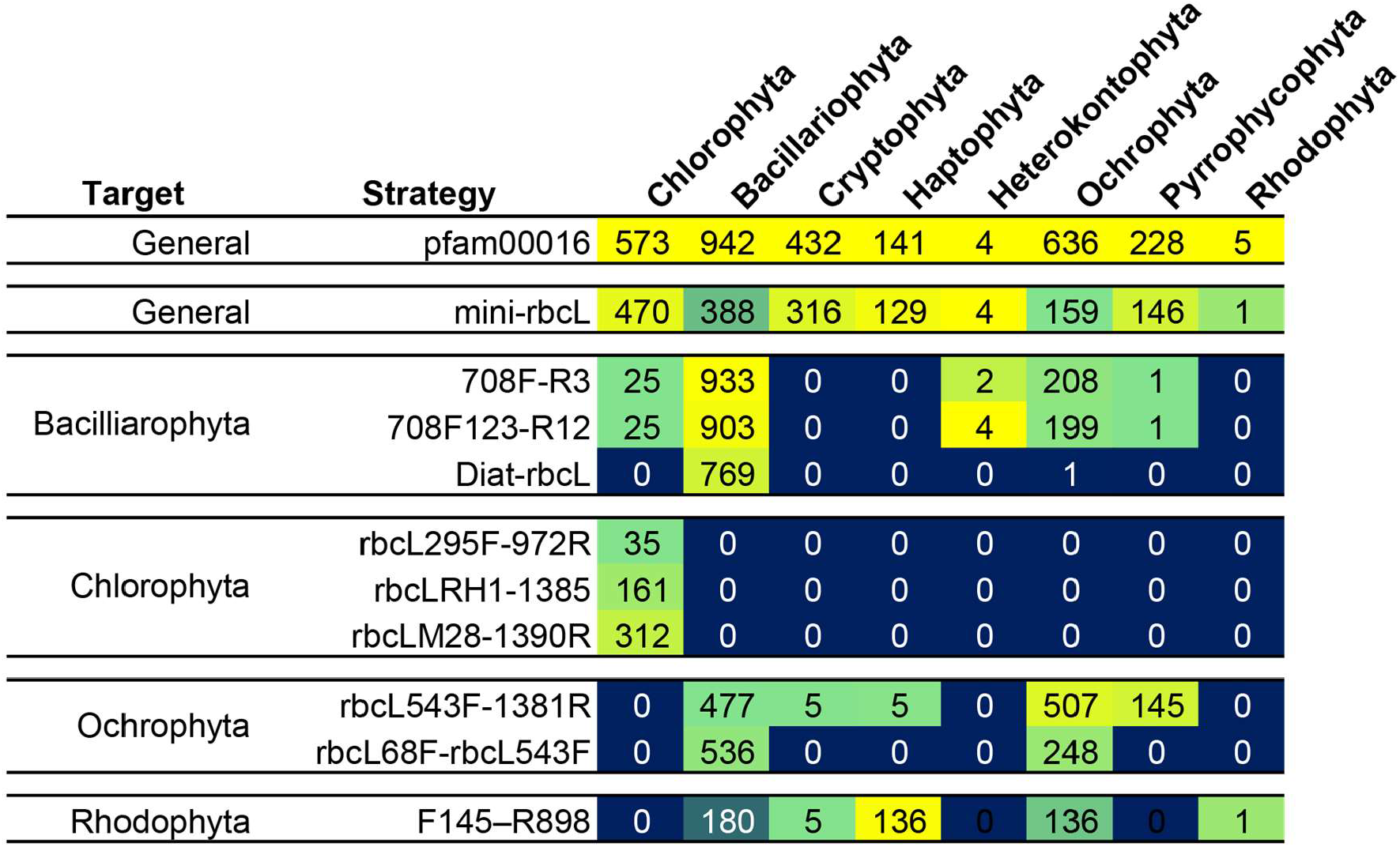
Success of *in silico* PCR applied to freshwater metagenome-derived *rbcL* sequences. The top row (pfam00016) represents the number of sequences derived from metagenome sequence searching, while all other rows indicate numbers of successful amplifications predicted using the indicated primer sets.

### WGS-based identification mirrors morphological identification

Until 2018, the NSF NEON program subjected freshwater aquatic samples to microscopic diatom and soft algae identification multiple times yearly. At the same sites, NEON continues to produce WGS metagenomes of benthic epilithic and epipsammic microbial communities. This offers a unique opportunity to directly benchmark the WGS *rbcL* classifier against classifications produced via expert examination of samples via microscopy (Figure 3). Microalgae representing 65 genera were identified by microscopy in 2018 at the West St. Louis Creek (WLOU) site. 22 of these genera did not have associated *rbcL* gene sequences in the BOLD database, meaning that they could not have been identified using sequencing tools. These 22 genera with no available *rbcL* sequences were quite rare in the community though, with an average relative abundance of 0.73% (16.1% of the community as a whole). Of the remaining 33 genera with available *rbcL* reference sequences, the *rbcL* WGS tool detected and correctly classified 10 and 16 in 2018 and 2023 metagenomes, respectfully. The remaining 23 and 17 genera were not detected in WGS data despite the classifier having been trained to detect them.

**Figure 3.**
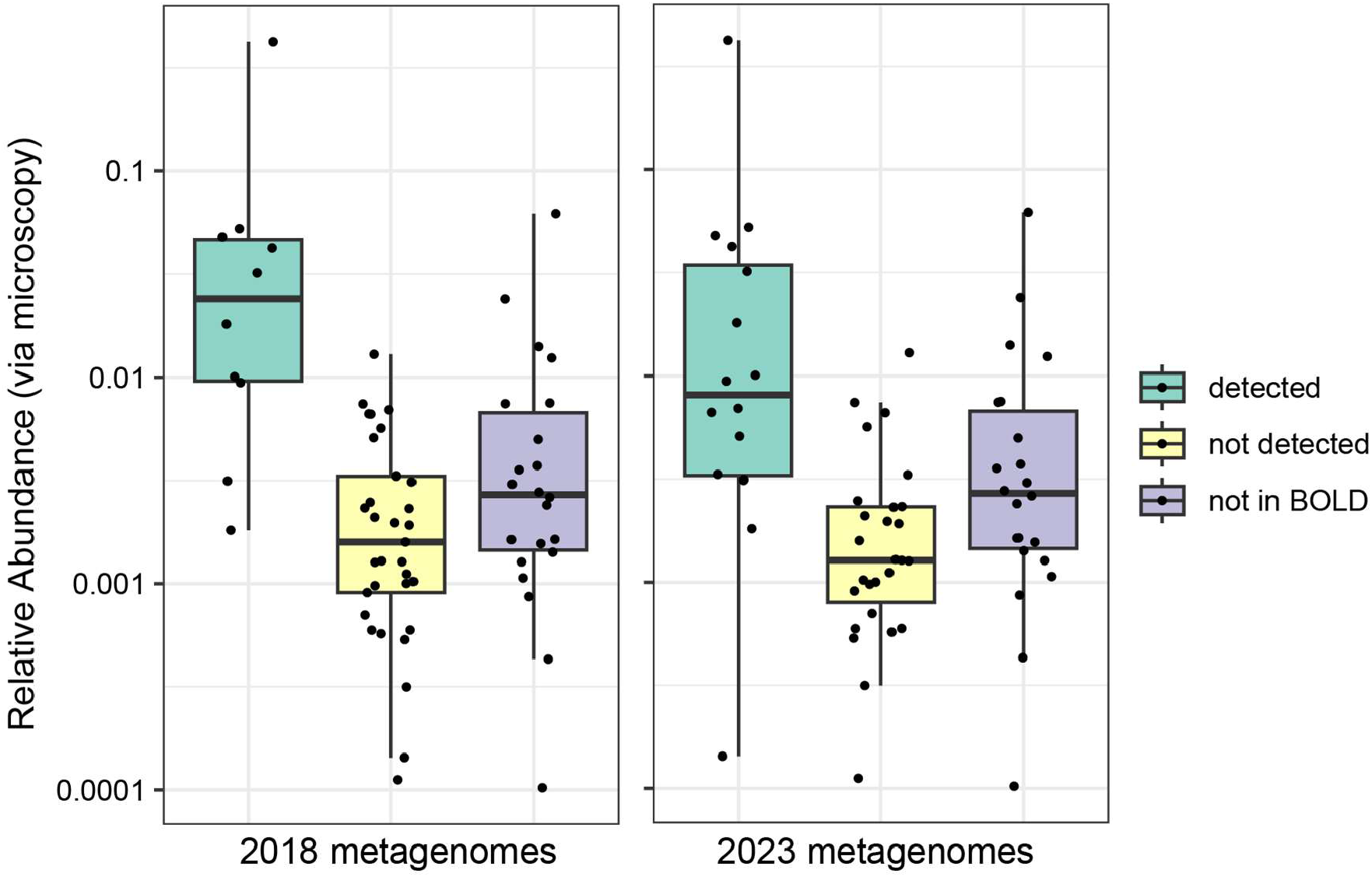
Benchmarking 2018 and 2023 metagenome-derived *rbcL* detections against 2018 microscopy-derived identifications at the NEON WLOU field site. Each point represents a microscopically-identified algal taxon. Points in the “detected” box plot were identified by metagenome analysis, points in “not detected” were not, while those “not in BOLD” represent algae with no available *rbcL* reference sequence. Relative abundances (y-axis) were calculated based on binned microscopic concentration data.

Genera that were detected by metagenome analysis were ∼20-fold more relatively abundant than those that were not detected. Mean relative abundances of the detected algae were 6.4% and 4.2% for 2018 and 2023 respectively, collectively representing 64.1% and 66.7% of the microalgal community. The non-detected taxa had mean relative abundances more than an order of magnitude lower, 0.27% and 0.23%. In other words, WGS-based detection was successful for more common taxa but failed to detect rare community members.

Ten genera were detected via WGS analysis that were not detected via microscopy, which presents the question as to whether these were false detections or whether metagenome-based detection offered an advantage over visual inspection. This included several microalgal genera that were could have been overlooked due to exceedingly small size or challenging morphology ^38^, including three of phylum Ochrophyta (*Ochromonas, Heterococcus*, and *Naegeliella*) and an exceedingly small lichen-symbiote Chlorophyte *Trebouxia*, only recently acknowledged as capable of a free-living lifestyle ^39^. Four macroalgae were also detected by sequence analysis, two Rhodophytes (*Batrachospermum* and *Audounella*) and the Chlorophytes *Ulva* and *Acrosiphonia*, which could be consistent with cellular debris, gametes, or spores which would be sequenced but not visually identified. Two diatoms were also detected by sequence analysis in only the 2023 samples, *Asterionella* and *Berkeleya. Asterionella* is common in freshwater systems around the world, and while *Berkeleya* is typically considered a marine taxon, it has been recently reported to be resident a freshwater river as well ^40^. Taken altogether, the algae that were identified by sequencing only are all plausible detections in freshwater systems, and most of them would likely be missed during routine microscopic identification, suggesting a possible strong advantage of WGS-based algal detection. However, without additional confirmatory empirical tests (ex. focused PCR assays) performed on environmental samples, determination as to whether these were authentic identifications is not possible.

### Case-Study: Simultaneous characterization of algae and prokaryotes allows for insights into bacterial-algal interactions

A recent study investigated the microbiological basis of methane production in Yellowstone Lake, a fully aerobic freshwater system ^41^. The study utilized 16S rRNA amplicon libraries to identify a responsible bacterial clade and, based on the metabolic potential of the organism, speculated that methylamines released from lysed algae may be the biochemical starting material. Eight metagenomes generated by the same investigators at that same site, five on the same day, were analyzed here to demonstrate the potential of simultaneous prokaryotic and algal taxonomic analyses for providing additional insights into such multi-domain phenomena. The single copy conserved protein (SCCP) ribosomal protein L2 was used to approximate the prokaryotic taxonomic community component in tandem with the *rbcL* classification pipeline, allowing for calculation of intra-sample bacterial and algal relative abundances.

The taxonomic analytic results indicated that while the dominant prokaryotes remained fairly consistent between sampling periods, the algal community showed strong variation in composition and overall relative abundance (RA) in the community across time and depth. The measured RA of the prokaryotic order Burkholderiales in July 24, 2017 samples (9.1 – 12.2%) corresponded well with measurements reported in the study (13%), indicating that the estimation of bacterial taxonomy by ribosomal protein L2 was appropriate. Cyanobacteria were dominant over algae the system, at 4.0 – 10.0% RA, vs. algae with 0.5 – 1.7% RA (although these numbers do not take into account marker gene copy numbers per cell). While the specific algae present in the system may offer insights into underlying drivers of methane production, the complete lack of algal detection in the sample corresponding to the highest methane spike (July 7, 2017 at 11.5 meters (Figure 4)) is consistent with the authors’ speculation that an algal die-off may have prompted the event.

**Figure 4.**
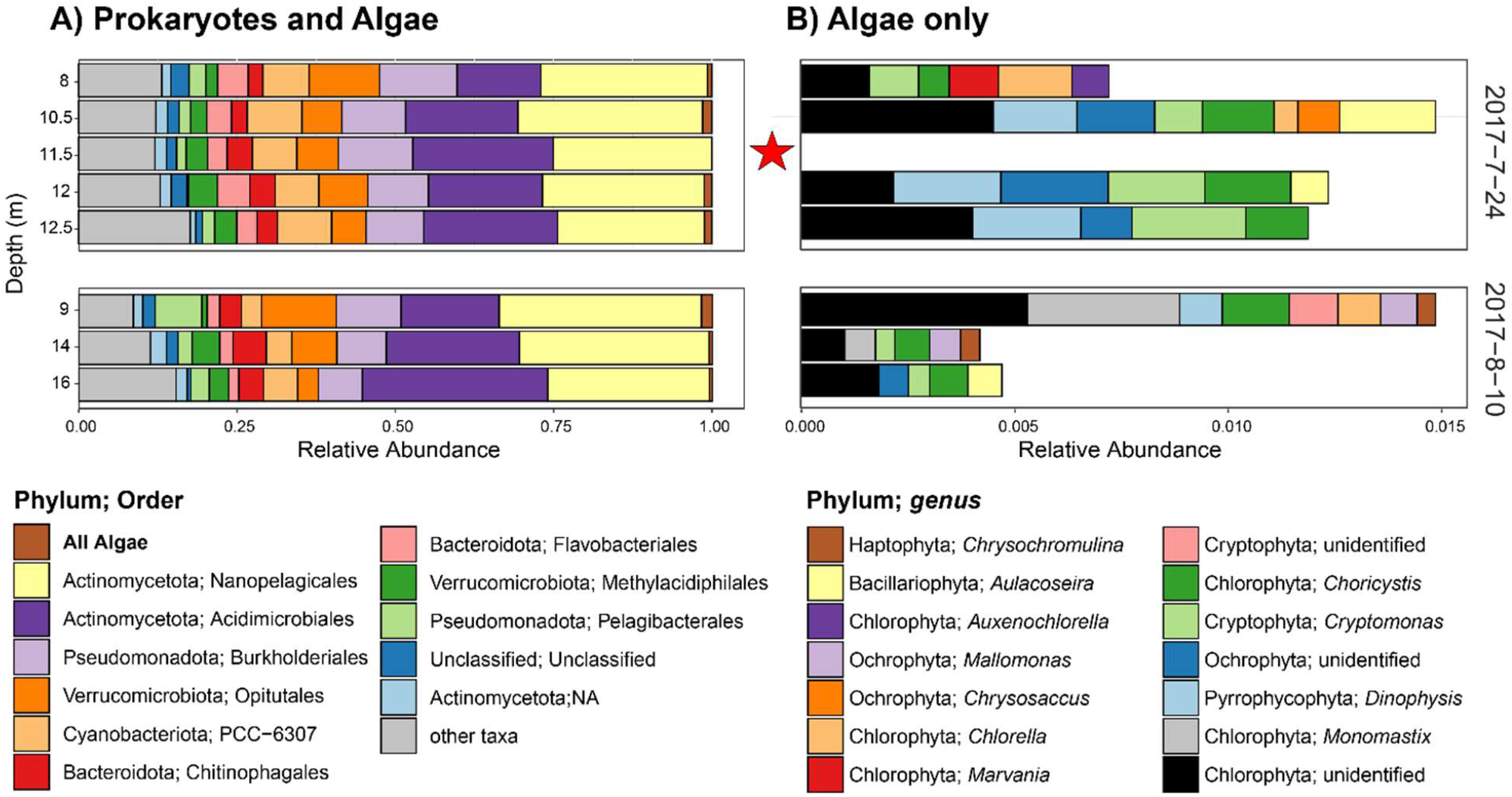
Metagenome-derived microbial community composition at Yellowstone Lake. Samples were collected on two different dates (shown on right) and at multiple depths (shown on left). The red star indicates the depth and timing of a methane spike as reported by Wang, et al. (2021). A) combined algae and prokaryotes, presented to order level taxonomy. B) Algae only, presented at genus-level taxonomy.

This case study demonstrates the insights that metagenomic-based broad spectrum algae detection can provide into complex environmental systems wherein the interactions between bacteria and algae drive important phenomena.

## Conclusions

Holistic assessment of diverse algal clades alongside prokaryotic communities has historically been hampered by several factors, especially by a reliance on taxonomic barcode PCR amplicon library sequence analysis. The rapidly decreasing costs of WGS metagenome sequencing is revolutionizing microbial community analysis. In this study, it was demonstrated that the algal barcode gene *rbcL* is efficiently detected by the conserved domain pfam00016 model. The AI/ML *rbcL* taxonomic classifier performed at near-perfect coverage and accuracy against a ∼48,000 sequence validation set down to the family or genus level for most algal clades and showed excellent performance to species level for many. The application of this model to *rbcL* sequences harvested from over 4,000 freshwater metagenomes generated a database of over 14,000 algal detections. Further, it was demonstrated by *in silico* PCR that available algal *rbcL* PCR primers likely fail to capture a significant proportion of these sequences, indicating that metagenome-based detection is more comprehensive than PCR.

A benchmarking of the metagenome-based tool against microscopy-derived identifications revealed contrasting advantages and disadvantages of the approach. Sequence-based detection successfully classified the more common algae. While less common algae were not detected, which is expected due to sequencing depth limitations, it was able to detect algal clades more recalcitrant to microscopic detection – the rapidly decreasing costs of sequencing will continue to make this less of an obstacle over time. A key limitation of the approach was revealed in that many of the microscopically identified algae have no available *rbcL* sequence.

Finally, a case study was presented whereby bacterial and algal community members were identified simultaneously, allowing for holistic calculations of sample-by-sample relative abundances. Analytic results were in accordance with previous findings in that particular environment and demonstrate the benefits of co-identification to achieve deeper understanding of important environmental phenomena. The interactions between prokaryotes and algae are likely extensive but poorly researched to date; this approach presents a methodology for future research. As sequencing costs continue to decrease and algal barcode gene sequences are deposited into databases, a metagenome-based approach for holistic freshwater community analysis will continue to improve. Furthermore, inclusion of classification models for other photosynthesizer barcodes such as *matK, trnH-psbA, trnL-F* will refine algal taxonomic precision, and eukaryotic barcode genes such as COX1 could extend the technique to include non-photosynthetic protists.

## Supplementary Materials

All supplementary materials are available for download at https://doi.org/10.6084/m9.figshare.29996962.

## Funding

This study was funded by Battelle Memorial Institute Internal Research & Development.

## Notes

https://doi.org/10.6084/m9.figshare.29996962

### Competing Interest Statement

The authors have declared no competing interest.

